# Non-invasive quantification of contractile dynamics in cardiac cells, spheroids and organs-on-a-chip using high frequency ultrasound

**DOI:** 10.1101/2022.06.28.497094

**Authors:** Eric M. Strohm, Neal Callaghan, Neda Latifi, Naimeh Rafatian, Shunsuke Funakoshi, Ian Fernandes, Milica Radisic, Gordon Keller, Michael C. Kolios, Craig A. Simmons

## Abstract

Cell-based models that mimic in vivo heart physiology are poised to make significant advances in cardiac disease modeling and drug discovery. In these systems, cardiomyocyte (CM) contractility is an important functional metric, but current measurement methods are inaccurate, low-throughput, or require complex set-ups. To address this need, we developed a standalone non-invasive, label-free ultrasound technique operating at 40-200 MHz to measure the beat rate, beat rhythm, and force of contraction of cardiac models, ranging from single adult CMs to 3D microtissue constructs in standard cell culture formats. The high temporal resolution of 1000 fps resolved the beat profile of single mouse CMs paced at up to 9 Hz, revealing limitations of lower speed optical based measurements to resolve beat kinetics or characterize aberrant beats. Coupling of ultrasound with traction force microscopy enabled the measurement of CM longitudinal modulus and facile estimation of adult mouse CM contractile forces of 2.34 ± 1.40 μN, comparable to more complex measurement techniques. Similarly, measurements of beat rate, rhythm, and drug responses of CM spheroid and microtissue models were demonstrated. In conclusion, ultrasound can be used for the rapid characterization of CM contractile function in a wide range of commonly-studied configurations ranging from single cells to 3D tissue constructs using standard well plates, with applications in cardiac drug discovery and cardiotoxicity evaluation.

## INTRODUCTION

Advances in induced pluripotent stem cell (iPSC) technologies have made it possible to create human cardiomyocyte (CM)-based models that mimic key aspects of the in vivo structure, function and physiology of heart tissue in health and disease [1]–[7]. This has driven significant interest in using 2D and 3D cell-based cardiac models to study and evaluate drug and stem-cell based therapies [8]–[10], with the unique possibility to directly evaluate contractile responses of human cardiac tissues.

However, measurement of contractility to evaluate CM function is not a routine in vitro assay. This is due, in part, to the complexity and/or deficiencies in current contractility measurement systems. The most common contractility tools include impedance plates, multielectrode arrays (MEAs), optical imaging methods, and custom techniques that require specialized substrates or methods. Impedance plates and MEAs [11] have good sensitivity and temporal resolution but require specialized substrates containing electrodes, are limited to 2D cultures, and cannot be used on single cells. Further, the measurements can be influenced by differences in confluence levels, cell connectivity, and impulse propagation, and cannot provide absolute contraction forces. Optical imaging can provide either high spatial resolution or high temporal resolution, but not both when imaging single CMs; clear optical access is also required. Other techniques to measure single CM contractility – including atomic force microscopy [12], bending posts [13], micropost arrays [14]–[16], thin film curling [17]–[19] or wrinkling [20], traction force microscopy (TFM) based on microbead movement [21], [22], and carbon nanotube strain sensors [23] – require specialized components or substrates and significant resources or skills, and tend to have low throughput, limiting widespread adoption. Options for assessing the contractility of 3D cardiac tissues are limited to either optical imaging [24], or custom systems such as two-post systems [25]–[27] including the Biowire II [28].

To address the deficiencies of current contractility measurement systems, we developed a non-invasive label-free technique to assess the beat rate, contraction force, and beat kinetics of single CMs and 3D myocardial microtissues using high frequency ultrasound. Clinical ultrasound (<15 MHz) is commonly used in cardiology for probing the heart size and shape, blood perfusion, and myocardial function [29]–[31], but does not have the spatial resolution to determine the beat characteristics of single cells and microtissues. High frequency ultrasound operating at frequencies over 200 MHz has high spatial resolution (<8 μm) to detect the signals from the top and bottom of the cell that enable evaluation of the mechanical properties at the single cell level [32]–[34]; however, limited pulse repetition rates and acquisition speeds have hampered its applications in rapid dynamic processes such as beating CMs. We developed a custom acoustic system to track transverse CM deformation during contraction using ultrasound frequencies from 80-200 MHz with capture rates up to 1 kHz, from which we estimate axial deformation and contraction force along the long axis of the cell. The 10 GS/s sampling rate can track the change in the propagation time of the ultrasound signals with 0.1 ns precision, translating to better than 100 nm length scales in the transverse direction. We demonstrate the technique on single adult mouse CMs, 3D hPSC-CM-based spheroids [35], and engineered CM tissue constructs using the Biowire II cell culture system [28]. As CM models and heart-on-a-chip systems mature and demonstrate advantages over animal models [36], versatile tools to measure important contractility metrics must capture the complex CM model dynamics. Our technique provides non-invasive contractile profiling on standard microtiter plates and custom fabricated devices, at unprecedented temporal resolution, and scales from single cells to microscale tissues by adjusting the ultrasound frequency. Optically clear devices are not required, as the ultrasound can penetrate opaque materials. This new measurement system will help improve understanding of CM contraction mechanics and drug efficacy and cardiotoxicity screening, ultimately leading to better cardiac therapeutic strategies.

## RESULTS

### Imaging system

Our custom designed ultrasound system was built on top of a commercial Olympus IX71 microscope to enable simultaneous optical and ultrasound imaging. A 200 MHz single element ultrasound transducer with a 0.5 mm focal length and an 8 μm spatial resolution was used for imaging single CMs. With a well plate positioned in the microscope, the ultrasound probe was positioned directly over the optical objective for precise positioning under optical guidance (Figure 1A,B). We designed the ultrasound pulse and acquisition hardware to enable rapid, precise pulse transmission and receiving. This was achieved by using a high digitization rate of 10 GS/s with a low pulse-pulse jitter of 30 ps. Contraction sequences were recorded at 1000 frames per second, where each frame was an average of a 50-pulse burst cycle performed at 500 kHz pulse repetition frequency within 0.1 ms, to increase the signal to noise ratio (SNR). The entire system was enclosed in an incubator maintaining 37°C.

**Figure 1:**
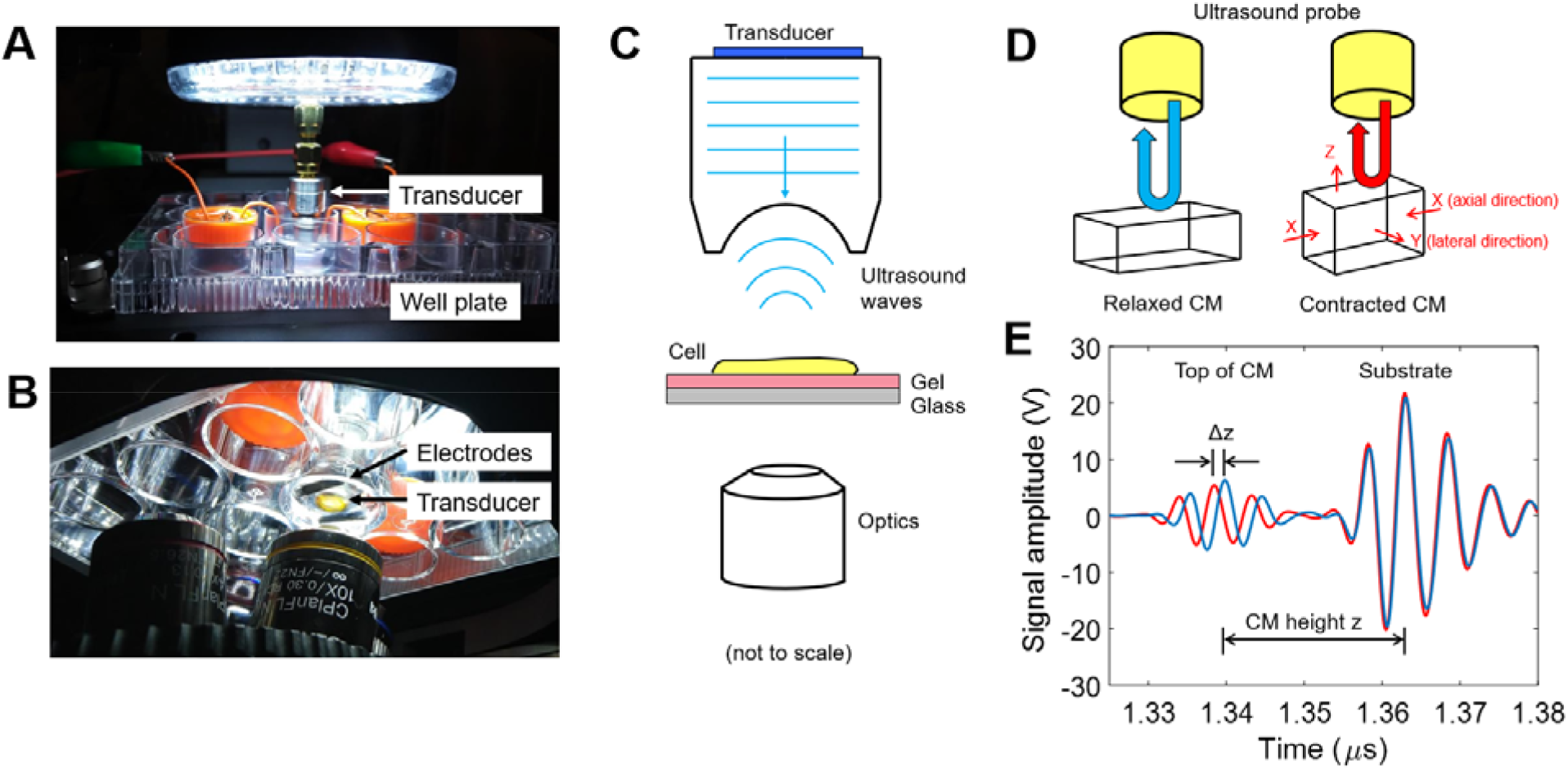
(A) Picture of the imaging system with the ultrasound probe immersed into the well plate. (B) A view from below, showing the ultrasound probe between the electrodes. (C) A schematic of the imaging system, with the cell located between the ultrasound probe and the optical objective. (D) When the CM contracts, its length decreases and its height increases. As the CM height increases, the time of propagation of the US wave decreases. (E) The ultrasound waves reflect from the top and bottom of the CM, with the reflections separated in time (blue curve). When the CM contracts and the height increases, the reflection from the top of the CM arrives sooner. The difference in the time of propagation between the relaxed and contracted states give the change in CM height, while the difference in the time of propagation between the top and bottom of the CM gives the cell height.

Cardiomyocytes were isolated from CD1 male mice using previously established methods [37], and then plated onto polyacrylamide gels of 11 kPa stiffness and containing 500 nm fluorescent microbeads in contraction-inhibiting medium to allow for CM adherence to the gel substrate. The well plate was then transferred to the imaging system, and the medium was replaced with modified Tyrode’s solution to enable contractions. Graphite electrodes were placed into the well plate, and the CMs were paced using 25 V/cm, 5 ms duration square pulses at a rate of 1 Hz (S48 Stimulator, Grass Instruments, USA). The ultrasound probe was immersed into the medium and positioned over contracting CMs, and then a 10 s contraction sequence was recorded using both ultrasound (at 1000 fps) and optical imaging (at 20 fps). Supplementary video 1 shows the change in the ultrasound signals during CM contraction synchronized with optical imaging. A schematic of the system set-up is shown in Figure 1C.

As the CMs were insonified with ultrasound waves, some of the wave energy would be reflected from the top of the CM back towards the transducer. The rest of the wave energy would travel through the CM, and then reflect from the cell-substrate interface. There was a time delay between these two waves as the second reflection takes longer to propagate back towards the transducer; the high ultrasound resolution (8 μm lateral and axial) and high sampling rate (10 GS/s) enables capturing the separation of these reflected waves with 100 ps precision (equating to 76 nm in distance in water at 37°C). The difference in the propagation time between the two reflected waves was used to calculate the CM height. As the CM contracted in the axial direction (i.e., along the long axis of the cell), the CM height would increase. As the CM height increased, the propagation time of the ultrasound wave reflected from the top of the CM decreased as it was closer to the transducer, while the propagation time from the bottom of the CM stayed the same (Figure 1D). The change in the CM height ΔZ was calculated from the change in propagation time Δ*t* from the CM surface using ΔZ = ½ *c*\*Δ*t*, where c is the sound speed in the medium at 37°C (1520 m/s). Then, the total CM height *Z* was found using the same equation, but with Δ*t* defined as the time delay between the CM top and bottom. This is depicted in Figure 1E, where two ultrasound waves are shown: blue indicates the ultrasound waves recorded during relaxation, and the red are at peak CM contraction. Optical images (brightfield or fluorescence) were recorded simultaneously at 20-24 fps (Retiga 2000R, QImaging, USA).

### Single cardiomyocyte contraction measurements

The beat profile and beat rate can be found by tracking the change in propagation time of the reflected waves from the CM surface as a function of acquisition time. Acquisition sequences were performed at 1000 fps for a 10 s duration, for a total of 10,000 frames. For each sequence, the change in cell height Δ*Z* for each frame was calculated using the relaxed CM state as a baseline (Figure 2A). This high temporal sequencing easily resolves the contraction dynamics and beat profile (Figure 2B). These measurements are done in the vertical direction directly above the CM, reflecting the contraction of the CM along its transverse axis. However, the axial contraction dynamics along the long axis of the CM are desired. Assuming the CM is rectangular in shape and incompressible, the change in length Δ*X* can be determined from the change in height Δ*Z* by

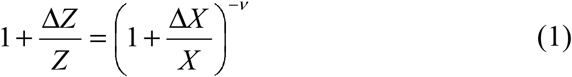

where *Z* is the CM height (measured with ultrasound), *X* is the known CM length (determined optically or by ultrasound imaging), and the Poisson’s ratio ν = 0.499, assuming near-incompressibility. This equation was solved for each Δ*Z* value from Figure 2A to obtain a plot of Δ*X* vs. time (Figure 2C). We recorded optical images at 20 fps simultaneously with the ultrasound measurements to compare the calculated length using ultrasound with the optical measurement (Figure 2D). While there was good agreement between the two methods for the majority of the 70 CMs measured (Figure 2E), ultrasound yielded significantly greater contraction of some cells (Figure 2E) and on average (Figure 2F; optical: 9.8 ± 4.1 μm vs. ultrasound: 11.5 ± 5.2 μm; p = 0.0007). This discrepancy may be partly due to assumptions inherent in our contraction model (Equation 1), including an idealized CM geometry and uniform boundary conditions, but as discussed below, it is likely explained mainly by the insufficient temporal resolution of 20 fps optical imaging that limits resolving the peak contractility fully.

**Figure 2:**
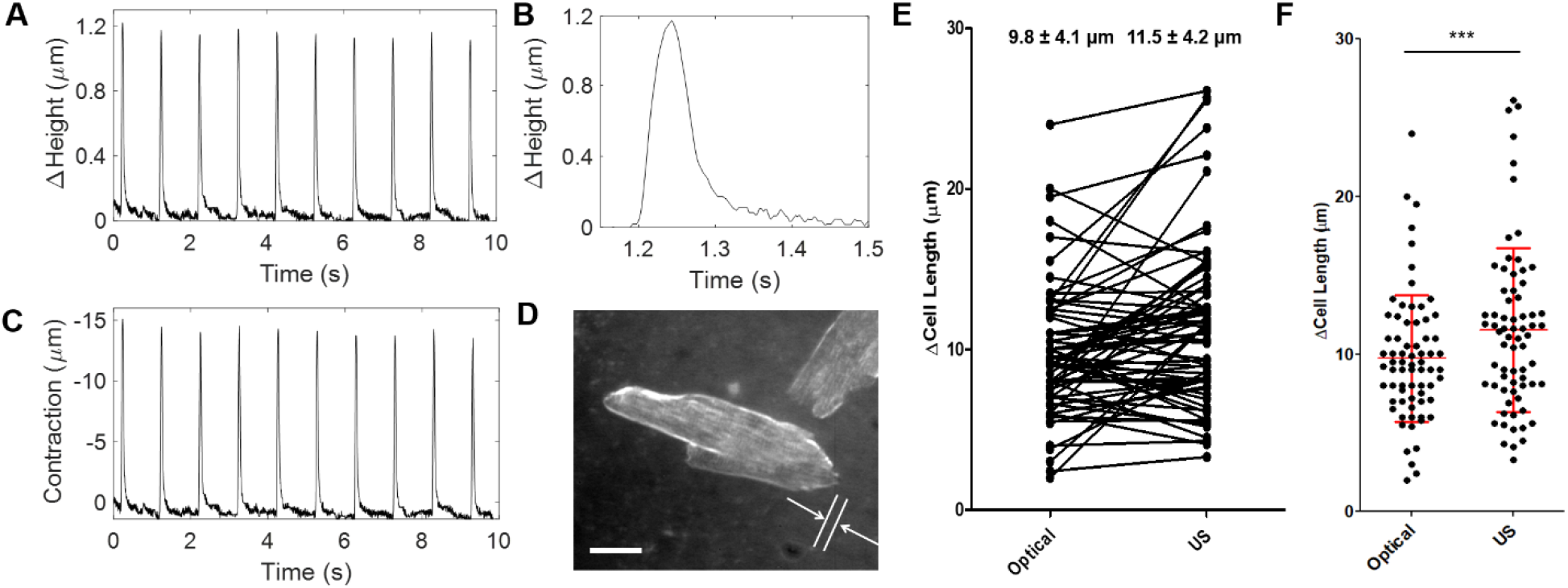
(A) The CM beat pattern is found by calculating the change in height throughout the acquisition time. (B) A single beat demonstrating the high temporal resolution. (C) The change in CM length (axial contraction), calculated assuming incompressibility and rectangular geometry. (D) The change in length was found using simultaneous optical imaging (scale bar 30 μm). (E) A pairwise and (F) boxplot comparison of the change in length between optical imaging and the ultrasound method (mean ± standard deviation, statistically different, *** p < 0.001).

To assess the temporal resolution of ultrasound imaging on the contraction measurement, contractions of a single mouse CM were measured when stimulated from 1 to 9 Hz (Figure 3A). As the pacing frequency increased, the change in cell height at peak contraction height decreased, from 0.7 μm at 1 Hz, to 0.38 μm at 5 Hz, to 0.2 μm at 9 Hz. Figure 3B shows a zoomed image where the beat cycle was 35 ms at 1 Hz, and 30 ms at 9 Hz (the beat cycle was measured at the full width at half maximum). To investigate the effects of the imaging frame rates on the estimation of contractile strain, a CM was measured using simultaneous ultrasound (at 1000 fps) and optical imaging (at 462 fps) using a uEye high speed camera (IDS, USA). Despite its higher frame rate, the optical sensitivity was lower and could not be used for fluorescent imaging at experimentally-relevant capture rates but was still sufficient to capture the CM edge movement during contraction using brightfield. The change in CM length was tracked using the Manual Tracking ImageJ plugin [38] by advancing through each image frame and manually placing markers on the CM edge during the contraction. A comparison of the ultrasound vs. optical method is shown in Figure 3C. At 462 fps, both the ultrasound and optical methods could clearly resolve the beat profile. This CM was paced at 1 Hz and the beat cycle lasted about 80 ms. The optical frame rate was decimated to 92 fps, and the beat shape could still be resolved. However, as the frame rate was decimated to 26 and 19 fps, only two data points were acquired during the beat cycle. The beat profile could not be resolved, and more importantly, the maximum contracted value could not be accurately determined. At 26 fps, the optical method measured a peak contraction that was 5-10% lower than the actual value; this discrepancy became more severe at 19 fps.

**Figure 3:**
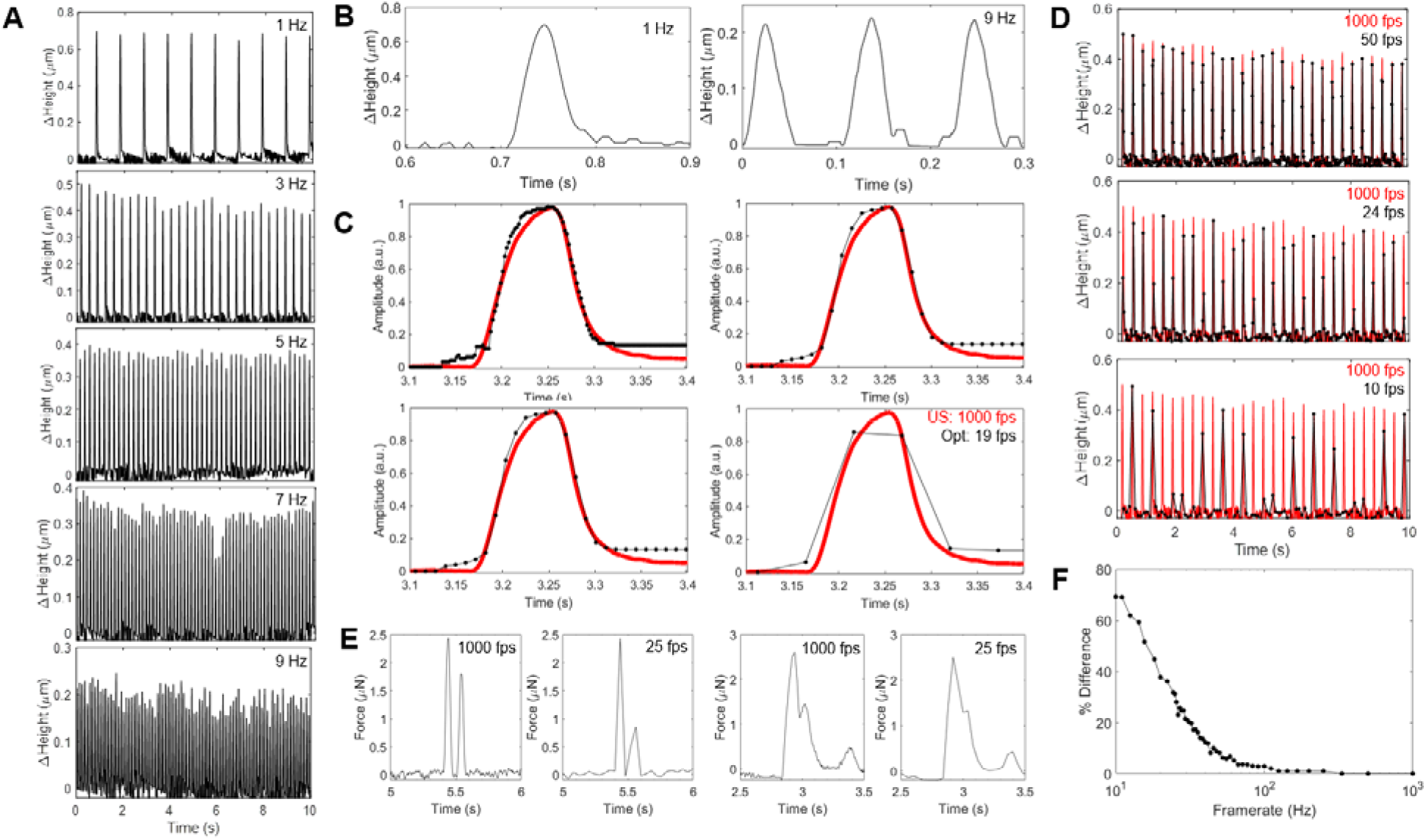
(A) The ultrasound system can acquire data at 1000 frames/sec, enabling high beat rate temporal resolution for CMs paced at up to 9 Hz. (B) A zoomed view of the beat rate at 1 and 9 Hz. (C) A single CM contraction measured simultaneously with ultrasound (at 1000 fps) and optical imaging (at 462 fps), paced at 1 Hz. The optical frame was decimated to 92, 24 and 19 fps to show how the reduced frame rate distorts the beat pattern, and limits the ability to measure the maximum contraction. (D) The 3 Hz plot in (A, red), was decimated to 50, 24, and 10 fps (black) to demonstrate how reduced framerates can miss the peak contractions. (E) The high frame rate can capture details such as close beats that become obscured when decimated to 25 fps. (F) The difference in peak contraction between 1000 fps and reduced frame rates (8% at 50 fps, 32% at 24 fps, and 69% at 10 fps).

The effect of the capture frame rate on the beat profile was examined by decimating the ultrasound-acquired beat pattern of the CM paced at 3 Hz from Figure 3A to 50, 24, and 10 fps. Figure 3D compares the original beat pattern captured at 1000 fps (red) to the decimated frame rate (black). At 50 fps, all of the beats were detected, but most were lower in amplitude than at 1000 fps; the average error between the beats measured at 1000 and 50 fps was 8%. At 24 fps, the error in amplitude was 32%, and then increased further to 69% at 10 fps. Many beats were missed at 10 fps, and at 24 fps, most beats had only one data point. To illustrate the effect that frame rate has on beat detection, two examples of beats very close together measured at 1000 fps with ultrasound, and then decimated to 25 fps, are shown in Figure 3E. The lower frame rate introduces errors in the beat shape, beat pattern, and beat separation, and in the second example, the second beat is nearly consumed within the first beat. The error between 1000 fps and lower framerates is shown in Figure 3F. The error in beat amplitude was less than 4% when frame rates above 66 fps were used, and less than 9% when frame rates above 42 fps were used. Significant errors were observed below 25 fps, where the error was greater than 25%. Comparing the CM length measurements using 20 fps optical imaging to the 1000 fps ultrasound imaging (Figure 2F), only 24% of the optical measurements were within 10% of the ultrasound values. The significant underestimation of peak contraction with low framerate optical imaging is consistent with our single cell observations (Figure 2E) and explains the observed discrepancies with ultrasound measurements (Figure 2F). The insight provided by the high resolution ultrasound measurements informs specifications for lower resolution methods: frame rates above 30 fps should be used to ensure beats are not missed and frame rates above 66 fps are needed for accurately resolved beat profiles.

### Single cardiomyocyte contractile force estimates

In addition to measuring beat rate, rhythm, and contraction strain, the force of contraction is a useful CM functional metric. The force of contraction, *F*, is often estimated from the contraction strain assuming linear elasticity according to

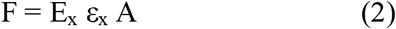

where *ε*_*x*_ is the axial strain, *A* is the cross-sectional area of the CM, and *E*_*x*_ is the elastic modulus in the axial direction. While strain and area can be determined by the ultrasound method described here, the elastic modulus cannot. We used atomic force microscopy (AFM) to determine the elastic modulus of single adult mouse CMs by nanoindentation, with an average value of 10.1 ± 2.2 kPa. However, cells have anisotropic mechanical properties, particularly CMs, which are elongated with sarcomeres and other structures positioned along the length of the cell [39]–[42]. AFM measurements of CMs typically assume isotropic properties based on measurement perpendicular to the axis of contraction, not the elastic modulus along the length of contraction.

To our knowledge, the only measurements of the axial elastic modulus of live CMs have been done on rat CMs subjected to tensile stretch, yielding equivalent moduli of ∼12 kPa, which was about six times stiffer than the transverse elastic modulus [43] However, these measurements may not reflect the moduli of a CM contracting auxotononically, which is arguably more physiological. To make those measurements with high temporal resolution, we combined the ultrasound method with traction force microscopy (TFM). TFM is a gold standard method to measure the contraction force of CMs cultured on soft gels embedded with nanoscale fluorescent beads that shift locally as a CM contracts (Figure 4A). The force exerted upon the gel by the CM can be calculated using the Fourier transform traction cytometry (FTTC) method [44], [45], based on the elastic modulus of the gel. Combined ultrasound-TFM measurements yielded the force (via TFM), strain (via ultrasound), and the cross-sectional area (width via optical or ultrasound imaging and height via ultrasound), from which the axial elastic modulus was estimated by Equation 2. For adult mouse CMs, we determined *E*_*x*_ = 46 ± 12 kPa, approximately four-fold greater than that measured by AFM nanoindentation, as expected and consistent with previous measurements [43]. The mean axial modulus was then used to calculate the contraction force of CMs that were measured using only ultrasound (Figure 2C). A representative plot of the force vs. time is shown in Figure 4B. In this example, the beat-to-beat peak force varied between 2.96 to 3.30 μN, with an average value of 3.10 ± 0.10 μN. This was repeated for 70 CMs, with an average force of 2.34 ± 1.40 μN (Figure 4C). These values, obtained for cells on standard culture substrates, are similar to what others have reported for adult mouse CMs using specialized culture formats and devices: 1.8 μN using traction force microscopy [46], 5.8 μN using a MEMS force gauge [47], 2-8 μN using a micro force gauge [48], 0.5-2 μN using a carbon fiber force-length control system [49], and 10 μN using magnetic bead movement [50].

**Figure 4:**
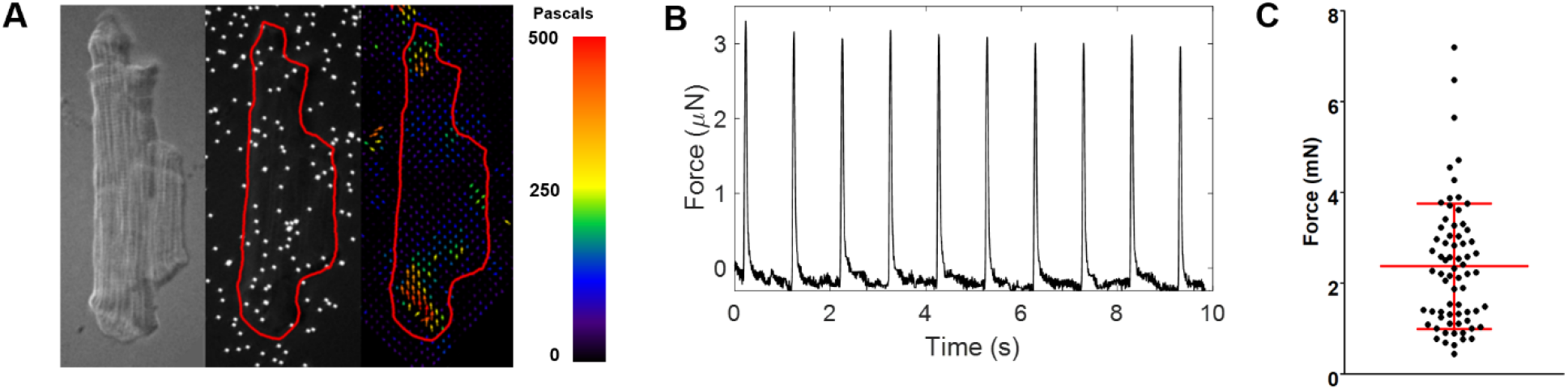
(A) Traction force microscopy (TFM) was used to determine the force of contraction of beating CMs. The CM axial elastic modulus was estimated from the force measured by TFM, and strain and cross-sectional area measured by ultrasound. (B) A typical contraction force curve for a CM, based on the axial elastic modulus estimate and contractility measured by ultrasound. (C) 70 CMs were measured using this technique, with an average force of contraction of 2.34 ± 1.40 μN (mean ± standard deviation).

### 3D microtissue measurements

A significant advantage of the ultrasound system is that it can examine different length scales, from single cells to mm-sized microtissue samples by changing the ultrasound frequency. 3D cardiac spheroids [35] 100-300 μm in diameter were examined using an 80 MHz ultrasound transducer, which had a 6 mm focal length, 25 μm resolution in the z-axis (direction of propagation (or) transverse CM direction), 60 μm resolution in the x-axis (perpendicular to the axis of propagation (or) or axial CM direction). The same measurement procedure described for the single CMs was used; the transducer was positioned above the spheroid, and ultrasound signals were acquired for 6 seconds. A representative optical image of the spheroid and corresponding ultrasound-acquired beat profile are shown in Figure 5A,B. The force calculation for the spheroids presents an additional difficulty, as the CMs within the spheroid are not aligned, which tends to result in irregular contractile behavior (supplementary video 2), which, in contrast to single CMs, cannot be accurately modeled using an idealized geometry estimate contractile force reliably. However, contractility can be measured with high temporal resolution to determine the beat rate and rhythm, which are important metrics for evaluating drug response. To demonstrate this, we performed a drug-dose curve using epinephrine (1-50 nM) and dofetilide (0.5-150 nM). As expected, the beat rate increased steadily with epinephrine treatment, with a final beat rate increase of 29% above baseline at 150 nM (Figure 5C). Similarly, for dofetilide, the beat rate increased steadily to peak at 53% above baseline at 50 nM (Figure 5C).

**Figure 5:**
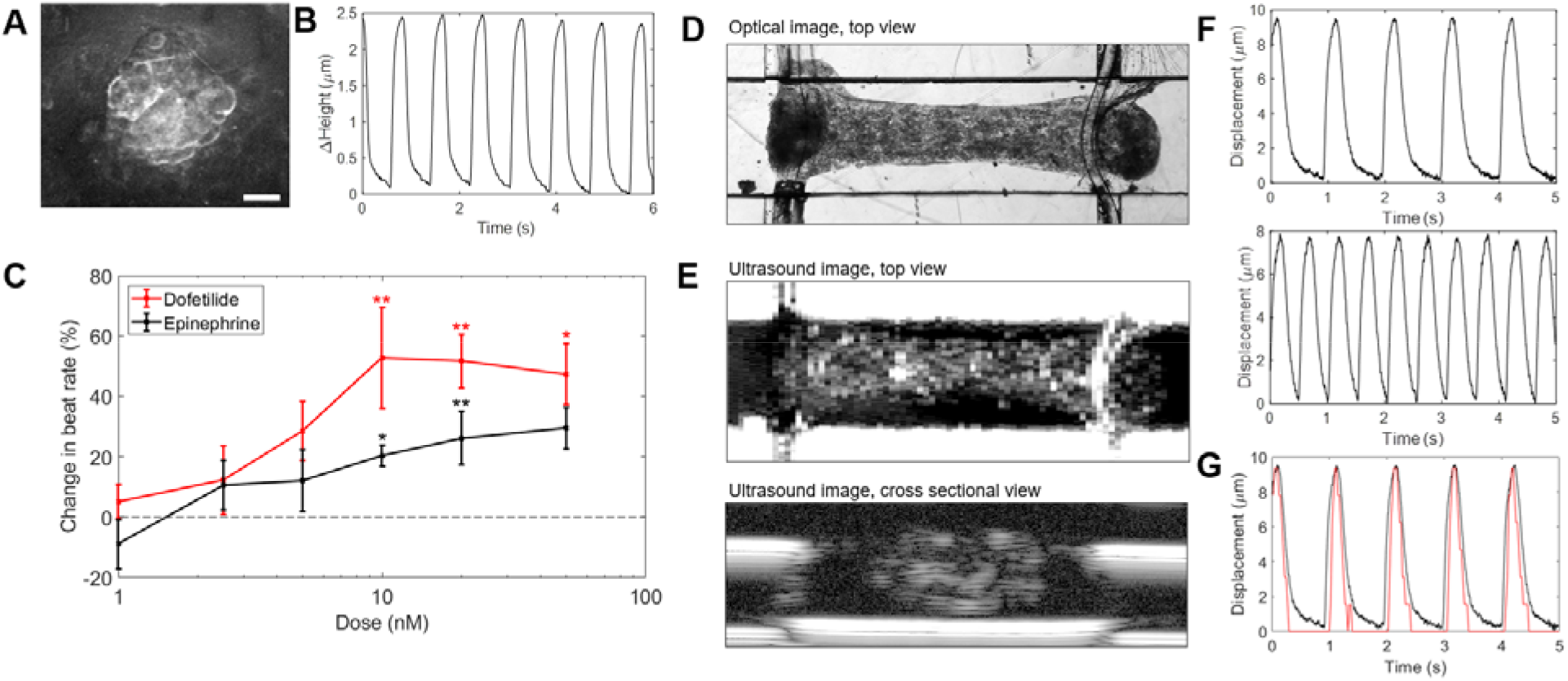
The ultrasound system can be used on larger tissue samples by changing the ultrasound frequency. (A) A representative spheroid 200 μm in diameter was measured using 80 MHz ultrasound, with the (B) beat pattern obtained using ultrasound. (C) Drug dose curves using epinephrine (n = 4 spheroids) and dofetilide (n = 5), mean ± standard deviation, * denotes P<0.05 relative to untreated control, dashed line denotes untreated control. In both cases, the beat rate increased with dose. (D) Optical image of the Biowire II. (E) Top and cross-sectional views of the Biowire II using 40 MHz ultrasound. (F) Ultrasound-measured beat pattern of the Biowire II stimulated at 1 and 2 Hz. (G) A comparison of the beat pattern of the Biowire II stimulated at 1 Hz using ultrasound (black) and optical tracking of the bending post (red).

The ultrasound system was then evaluated using the Biowire II, a 3D microtissue platform used to mature cardiac tissues and evaluate drug toxicity [51]. In this system, iPSC-CMs are suspended between two wires, which deflect when the CMs contract (Figure 5D). The deflection is measured optically to determine the beat rate and force of contraction. Using the same ultrasound system as above, the Biowire was scanned to create an overhead view (ultrasound C-scan) and a cross sectional view through the centre (ultrasound B-scan) to determine the Biowire shape and dimensions (Figure 5E). The ultrasound probe was then positioned over the middle of the Biowire, and ultrasound measurements were performed using stimulation at 1 and2 Hz, using the sample procedure as with single CMs. The ultrasound technique could detect the top and bottom of the Biowire, but no change in width or height was detected; the Biowire shifted with each contraction. These observations suggest that, unlike single CMs, the Biowire tissues compress under contraction. This is likely due to the densification of regional heterogeneity in the cellularity and extracellular matrix throughout the tissue volume, as observed in the cross-sectional ultrasound images (Figure 5E). Regardless, the movement measured by ultrasound was sufficient to determine beat rate and profile, which was similar to that of the CM spheroids, with rapid shortening and a slightly slower relaxation when stimulated at 1 or 2 Hz (Figure 5F). Of note, while the beat rates determined by ultrasound and 20 fps optical imaging were identical, the temporal resolution of the standard optical method was too low to resolve the fine details of the beat profile that were observed with ultrasound (Figure 5G). Because of its high temporal resolution and ability for label-free, non-invasive cross-sectional imaging (Figure 5E), the ultrasound system could also provide insights into local mechanics of beating 3D microtissues that are difficult to probe otherwise.

## Discussion

We have demonstrated that high frequency ultrasound imaging can detect and resolve the contractile profiles of single CM cells and microtissues up to several hundred micrometers in size. For single CMs, the estimated forces of contraction were comparable to other more complex techniques that required invasive probes or specialized substrates. These measurements can be done on CMs, organoids, or microtissues cultured on any type of substrate; in this study, 11 kPa polyacrylamide gels were used so that TFM measurements could be done simultaneously, but we have also measured CM contraction by ultrasound on standard polystyrene plastic cell culture plates, glass slides, and polyacrylamide with a stiffness as low as 2 kPa. Similarly, the technique is easily adaptable to custom culture plates, devices, and even cardiac-on-a-chip systems. It does not require researchers to adapt their assays to a specific substrate or material.

And while the measurements here were done with a sterilized ultrasound probe immersed into the cell media, it is possible to invert the transducer to perform the measurements from below the culture plate, in which case the thickness of the culture plate will dictate the requirements for the focal length of the ultrasound probe.

The ultrasound method can measure CM beat profile with temporal resolution (1000 fps here) that is unmatched compared to optical or multi-electrode array systems. In electrophysiology, the beat shape and profile can provide information about the contraction dynamics that can be used to evaluate pathophysiology, and drug effects but do not allow for absolute contractility measurements. In our system, the beat profile is based on the physical movement as the CM contracts, which, as demonstrated in Figure 3E, may allow detection of movement irregularities that can occur in arrythmias or other disorders that cannot be observed using electrophysiology. Future work will compare similarities contraction and ion transport similarities, which may provide new insight into the relationship between signaling and the contraction mechanics.

By coupling the ultrasound method with TFM, we were uniquely able to estimate the axial elastic properties of individual CMs, enabling contraction force estimates that were comparable to those measured with more complex and specialized set-ups. However, elastic properties vary from cell to cell, change depending on the degree of CM contraction, and are heterogeneous within organoids and microtissues [43], [52], [53]. This complexity cannot be comprehensively captured using the methods here. However, high frequency ultrasound may have unique advantages in addressing this limitation, as the system could be adapted in the future for high resolution shear wave elastography [54], [55] to measure the cell and tissue elastic properties needed for direct contraction force calculation.

## Conclusions

The ultrasound system detailed here represents a non-invasive, non-destructive, and label-free means by which to characterize cardiac cell or tissue contractility and mechanics in vitro. It is highly versatile and applicable to various culture formats and spatial scales, from single cells to complex 3D microtissues. Furthermore, this technique could be multiplexed in real time with other in situ monitoring (e.g., Ca^2+^ or voltage fluorescence imaging), or even complexed with optically opaque modalities (e.g., multi-electrode arrays) for high-yield physiological experimentation. The high spatial and temporal resolution allows for a novel degree of monitoring of live cell behaviour in physiologically-relevant mechanical environments. By providing new insight into myocardial contractile kinetics and beat profile, this system may allow for enhanced characterization of the mechanical aspects of pathophysiology and drug responses, therefore contributing to higher-value in vitro experimentation, personalized medicine, and drug screening applications.

## Materials and Methods

### Cardiomyocyte culture

Primary adult cardiomyocytes were isolated by previously-established methods from 8 week old CD1 male mice [37]. Cells were plated onto 250 μm thick, 11 kPa polyacrylamide gels as described previously [46], except that Geltrex was mixed into the pre-polymerized gel solution at a final concentration of 16.7%. Gels were cast onto 18 mm round, APTES-functionalized glass coverslips and embedded with 500 nm fluorescent nanobeads on their top surface. To inhibit contractions, cells were incubated with Tyrode’s solution with 15 μmol L-1 blebbistatin (Toronto Research Chemicals, Canada) [46]. Prior to measurements, cells were induced to contract spontaneously using sterile modified Tyrode’s solution containing (in mmol L^-1^) NaCl (130), KCl (5), NaH_2_PO_4_ (0.5), D-glucose (10), HEPES (10), taurine (10), MgSO_4_ (1), and CaCl_2_ (1.8), pH 7.4, as well as 1X chemically defined lipid concentrate (11905031, Gibco/Thermo Fisher Scientific).

### Biowire fabrication

Myocytes were differentiated from the human induced pluripotent stem cell (hiPSC) line BJD1 (gift from Dr. William Stanford, now at Ottawa Hospital Research Institute) following established monolayer differentiation protocols [56], [57]. Dissociated hiIPSC derived myocytes were mixed with human cardiac fibroblasts (Lonza, NHCF-V) at the ratio of 9:1. The cell mixture was resuspended in a collagen/Matrigel based hydrogel at the concentration of 3 mg/mL and 150 µL/mL, respectively. The hydrogel containing cells was then seeded in the BiowireII platform [51]. Throughout the first week of culture, the microtissue compacted and formed around the wires, suspending the microtissue.

### Spheroid fabrication

For ventricular cardiomyocyte differentiation for cardiac spheroids, a modified version of the embryoid body (EB)-based protocol was followed [58]. hPSC populations (HES2) were dissociated into single cells (TrypLE, ThermoFisher) and re-aggregated to form EBs in StemPro-34 media (ThermoFisher) containing penicillin/streptomycin (1%, ThermoFisher), L-glutamine (2□mM, ThermoFisher), transferrin (150□mg/ml, ROCHE), ascorbic acid (50□mg/ml, Sigma), and monothioglycerol (50□mg/ml, Sigma), ROCK inhibitor Y-27632 (10□uM, TOCRIS) and rhBMP4 (1□ng/ml, R&D) for 18h on an orbital shaker (70□pm). On day 1, the EBs were transferred to mesoderm induction media consisting of StemPro-34 with the above supplements, excluding ROCK inhibitor Y-27632 and rhBMP4 (8□ng/ml), rhActivinA (12□ng/ml, R&D) and rhbFGF (5□ng/ml, R&D). On day 3, the EBs were harvested, dissociated into single cells (TrypLE), and re-aggregated in cardiac mesoderm specification media consisting of StemPro-34, the Wnt inhibitor IWP2 (2□uM, TOCRIS) and rhVEGF (10□ng/mL, R&D). On day 5, the EBs were transferred to StemPro-34 with rhVEGF (5□ng/ml) for another five days and then to DMEM high glucose (4.5□g/l, ThermoFisher) media with compaction factors (CHIR (1□uM, TOCRIS), IGF2 (25□ng/ml, R&D)) and human insulin (10□ng/ml, Sigma) at day 10 for another six days. From day 16 to day 18, the EBs were transferred to DMEM high glucose media with XAV (4□uM, TOCRIS) and then transferred to maturation media [DMEM containing low glucose (2□g/L) with Palmitic acid (200□uM, Sigma), Dexamethasone (100□ng/ml, Bioshop), T3 hormone (4□nM, Sigma) and GW7647 (PPARA agonist, 1□uM, Sigma] for the following nine days. Finally, the EBs were cultured in DMEM containing low glucose supplemented with palmitic acid (200□uM) alone for the following five days (a total of 32 days). Cultures were incubated in a low oxygen environment (5% CO2, 5% O2, 90% N2) for the first ten days and a normoxic environment (5% CO2, 20% O2) for the following 22 days. From day 10 to day 32, the EBs were cultured in polyheme-coated low binding 10□cm culture dishes on an orbital shaker (70□pm).

### Ultrasound system

A custom ultrasound system was designed and built to rapidly insonify and acquire ultrasound signals with high precision. An Intel i7 computer with a trigger card (Spincore, USA) was used to control the hardware. A pulse generator (Geozondas, Lithuania) generated monocycles pulses that were sent through an RF-switch (Mini-Circuits, USA) to an ultrasound transducer (200 MHz transducer for single CMs, 80 MHz transducer for microtissues). Returning signals were amplified by a 30 dB amplifier (Miteq, USA) before digitization at 10 GS/s with 14-bit resolution (Teledyne SP Devices, Sweden). Cells were insonified at 1000 samples/sec, where each sample was the average of a 50-pulse burst with a 500 kHz pulse repetition frequency. The 50-pulse burst was used to increase the SNR, and occurred within 0.1 ms; we assumed there was negligible cell movement during this time. A 3-axis stage (Thorlabs, USA) was mounted to an Olympus IX71 microscope stage to move the ultrasound transducer over each individual cell under optical guidance (fig. 1). A total of 10,000 frames (10 s acquisition time) were then acquired with simultaneous optical video using a QImaging Retiga 2000 at 20-24 fps, or an IDS UI-3140CP high speed camera at 462 fps. The entire microscope system was enclosed in a temperature controlled incubator with a temperature setpoint of 37°C (In Vivo Scientific, USA).

### Contraction measurements and force estimates

CM contraction was determined by relating the change in height of the CM during each contraction (measured using ultrasound) to an axial deformation by assuming a rectangular prism geometry and uniform, incompressible contraction (eq 1). First, the baseline propagation time of the ultrasound echo from the cell surface at relaxation (e.g., 1.340 μs in Figure 1E) was found. As the CM contracts, the cell height increases (e.g., to 1.339 μs in Figure 1E); the change in CM height was calculated for each frame using the equation

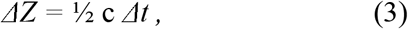

where *c* is the sound speed in the liquid (1520 m/s) and Δ*t* is the difference in propagation time between the baseline relaxation time and the signal at each frame. The transverse strain ε_z_ = Δ*Z*/*Z* was calculated using the change in cell height Δ*Z* and the cell height *Z*, measured at relaxation. The axial strain ε_x_ in the direction of contraction was then calculated using Poisson’s ratio ν = dε_x_/dε_z_, where ν = 0.499 for a nearly-incompressible cell, assuming rectangular prism geometry (Equation 1). The stress σ_x_ was then calculated using the linear elastic equation

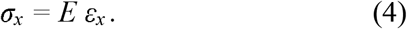

where E is the elastic modulus. This equation assumes that the CM is a homogeneous, isotropic, incompressible, elastic material. The longitudinal force exerted by the cell was then calculated using

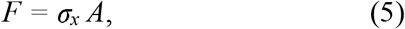

where *A* is the cross-sectional area of the cell, with the cell width obtained through optical imaging and the cell height obtained through ultrasound. These calculations were performed for every successive signal acquired, resulting in plots of Δheight, Δlength, and force measurements vs. time.

### Contractile force measurement using traction force microscopy

Fluorescent polystyrene microbeads 500 nm in diameter (Polysciences Inc, USA) were embedded in polyacrylamide gels with a stiffness of 11 kPa. Videos of beating CMs and the fluorescent beads were acquired at a framerate of 20-24 fps using an Olympus IX71 microscope with a QImaging Retiga 2000 CCD camera and Micromanager 1.4 [59]. The displacement field was measured by tracking the motion of the fluorescent beads. An iterative particle image velocimetry (PIV) method was used, where the image was divided into increasingly smaller interrogation windows to determine the displacement vectors. The traction field was then calculated using the Fourier transform traction cytometry (FTTC) method using ImageJ plugins [44], [45]. FTTC-determined stresses were integrated within the cell area to give a total cell force scalar, given the assumptions of uniaxial contraction and zero vector sum [60].

### Elastic modulus estimation

A JPK atomic force microscope (AFM; Bruker JPK NanoWizard 4, Cambridge, United Kingdom) was used to determine the elastic modulus of the polyacrylamide gels and the elastic modulus of non-beating adult CMs in the transverse direction. The indentation tests were performed using force spectroscopy contact mode at room temperature. Tip-less silicon nitride AFM cantilevers (Bruker, MLCT□O10, cantilever D with nominal spring constant of 0.03 N/m) were functionalized using 10 µm radius spherical polystyrene beads (Phosphorex Inc. Hopkinton, MA). A contact-based thermal tune method was applied to determine the precise spring constants. Each hydrogel sample was indented in three locations composed of a 10□μm□×□10□μm area in which four points were indented. The indentation tests were repeated five times at every indentation point, i.e., five technical replicates per point. For the adult CMs, cells were indented at three locations in their center and three locations at the cell end. Force-deflection curves were recorded, and the elastic modulus was obtained from the extend curves using the Hertz/Sneddon model. Data analysis was performed using the JPK Data Processing software (version 6.3.11). CM cells and tissues are strongly anisotropic [40]–[42]. AFM measurements yield the elastic modulus of the CMs along the transverse direction, perpendicular to the axis of contraction, while our linear elastic equation (4) requires the axial stiffness, which is unknown for single contracting CMs. Finally, the elastic modulus may be a dynamically changing parameter during contraction. For these reasons, we decided to use an experimentally extracted elastic modulus that was calculated using a combined ultrasound-TFM method. Equations 4 and 5 were combined and isolated to solve for *E*,

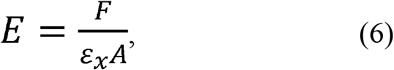

where the force (*F*) was measured using TFM, the cross-sectional area (A) using optical imaging (width) and ultrasound (height) assuming rectangular cross-section, and the axial strain (*ε*_*x*_) using ultrasound. The value of E, based on measurement of n = 6 CMs, was used in the ultrasound force determination calculations using equation 4.

## Acknowledgements

We are grateful to the Collaborative Advanced Microscopy Laboratories (CAMiLoD) at the University of Toronto for the use of their atomic force microscope. This study was funded by the Medicine by Design New Ideas Fund (MbDNI-2017-01) and a Collaborative Health Research Program grant from the Natural Science and Engineering Research Council of Canada (NSERC; CHRPJ 5083) and Canadian Institutes of Health Research (CPG-151946). ES was supported by an NSERC postdoctoral scholarship and a Medicine by Design Postdoctoral Fellowship. NIC was supported by a C. David Naylor Fellowship Endowed by a Gift from the Arthur L. Irving Foundation, Ontario Graduate Scholarship, NSERC Vanier Canada Graduate Scholarship, and NSERC CREATE TOeP Scholarship. NL was supported by a NSERC postdoctoral scholarship and a Fonds de recherche du Quebec–Nature et technologies postdoctoral fellowship award.

